# Human cytomegalovirus triggered necroptosis is suppressed by sequestration of MLKL in the nucleus of infected monocytes

**DOI:** 10.1101/2025.04.28.651015

**Authors:** Brittany W. Geiler, Shima Moradpour, Ben B. Chauder, Dilruba Akter, Gary C. Chan

**Affiliations:** Department of Microbiology & Immunology, SUNY Upstate Medical University; Syracuse, NY 13210, USA

**Author notes:** Corresponding author. Gary C. Chan. Ph.D., Professor, Department of Microbiology and Immunology, SUNY Upstate Medical University, 750 East Adams Street, Syracuse, NY 13210.

## Abstract

The systemic spread of human cytomegalovirus (HCMV) is associated with severe morbidity and mortality in immunocompromised and immunonaïve patients. Hematogenous dissemination of HCMV to different organ sites is facilitated by peripheral blood monocytes. Circulating monocytes have a short lifespan due, in part, to their intrinsic biological programming to initiate caspase 8-mediated apoptosis upon entry into the circulation from the bone marrow. We previously reported that HCMV extends the lifespan of infected monocytes by blocking procaspase 8 cleavage, yet the precise viral mechanism responsible for suppressing caspase 8 activity remains unknown. Here, we demonstrate that HCMV entry into monocytes rapidly increases the abundance of the antiapoptotic cellular FLICE-like inhibitory protein long (cFLIP_L_), which prevents procaspase 8 cleavage into active caspase 8. However, others have demonstrated that inhibition of caspase 8 opens a “trapdoor” cell death response termed necroptosis. Accordingly, we found the increased levels of cFLIP_L_, along with a co-stimulatory signal from toll like receptor 3 (TLR3), activates the receptor-interacting protein kinase 3 (RIPK3) responsible for initiating necroptosis. Despite triggering of the necroptotic cascade within infected monocytes, the final execution of this death pathway is thwarted by nuclear sequestering of mixed lineage kinase domain like pseudokinase (MLKL), the executioner of necroptosis. Together, our data reveal a multitude of countermeasures employed by HCMV to obstruct cellular antiviral death responses within infected monocytes.

**Importance:** HCMV is highly prevalent in the adult population with a seroprevalence of 50-80% in the United States. Although immunocompetent individuals are generally asymptomatic, HCMV infection can cause multiorgan disease in immunocompromised and immunonaïve patients. Peripheral blood monocytes are responsible for the systemic dissemination of HCMV. However, the inherently short lifespan of monocytes combined with the induction of antiviral cellular death responses requires HCMV to circumvents cell death pathways to allow for viral spread. In this work, we show that HCMV induces cFLIP_L_ levels to inhibit caspase 8-mediated apoptosis. However, the inhibition of apoptosis, combined with TLR3 activation, triggers a secondary cell death pathway termed necroptosis. As a countermeasure to block necroptosis, HCMV sequesters MLKL within the nucleus of infected monocytes. Defining the precise mechanisms through which HCMV stimulates survival will provide insight into novel therapeutics able to target infected monocytes.

## Introduction

Human cytomegalovirus (HCMV) is highly prevalent in the adult population with a seroprevalence of 40-100% globally (1–3). In immunocompetent individuals, HCMV infection is generally asymptomatic, although HCMV can cause acute infectious mononucleosis (4). HCMV has also been linked to the development of chronic inflammatory diseases, such as atherosclerosis and restenosis, as well as cancers, including glioblastoma and breast cancer (5–11). In contrast to individuals with fully matured immune systems, more than 40,000 immunonaïve neonates are born with congenital HCMV each year, resulting in upwards of 8,000 children with permanent hearing, vision, or neurological deficits (2, 12–14). In immunocompromised individuals, including AIDS patients and transplant recipients, HCMV infection is a significant cause of organ pathologies, which are characterized by widespread viral dissemination and inflammation leading to end-organ damage (15, 16).

During an acute infection, HCMV utilizes peripheral blood monocytes to systemically disseminate from the initial point of infection. HCMV stimulates the survival and differentiation of naturally short-lived monocytes non-permissive (quiescent) for viral replication (17–19) into long-lived macrophages fully permissive for replication (20–22). In order to promote the differentiation of infected monocytes, HCMV opposes the biological programming of monocytes to rapidly initiate both intrinsic and extrinsic apoptosis upon exit from the bone marrow, which can be further accelerated by DNA damage, reactive oxygen species, or pathogen infection (23–25). Intrinsic apoptosis is triggered by permeabilization of the mitochondria membrane, release of cytochrome C, and subsequent activation of executioner caspases such as caspase 3 (25, 26). We previously reported that HCMV stimulates the increased production of a select subset of antiapoptotic proteins responsible for arresting multiple steps of the intrinsic apoptotic pathway (16, 22, 27). Conversely, extrinsic apoptosis is triggered by cell death receptors that activate initiator caspases, including caspases 8 and 10, to directly cleave and activate executioner caspases (28, 29). HCMV abrogates the cleavage of procaspase 8 into activated cleaved caspase 8 within infected monocytes halting the progression of the extrinsic apoptotic pathway (22, 30). Although the HCMV *UL36*-encoded viral inhibitor of caspase 8 activation (vICA) directly blocks the cleavage of procaspase 8 (31, 32), its lack of expression during quiescent infection of monocytes indicates a distinct yet to be identified mechanism through which HCMV suppresses extrinsic apoptosis.

When extrinsic apoptosis is blocked, a “trapdoor” cellular death pathway termed necroptosis is opened (33). Necroptosis is a caspase-independent programmed cell death pathway that requires two conditions for activation: the inactivation of caspase 8 and an initiating signal from a cell death receptor (34–37). Canonical necroptosis signaling is initiated through TNFR1 via the recruitment of TNFR1-associated death domain (TRADD) and receptor-interacting serine/threonine-protein kinase 1 (RIPK1) to the cell membrane (38, 39). RIPK1 is then deubiquitinated and forms complex IIa with TRADD and FAS-associated via death domain protein (FADD) (38). The formation of complex IIa is a pivotal point whereby extrinsic apoptosis is triggered by recruiting and cleaving procaspase 8 (40). However, the presence of inhibitors of procaspase 8 cleavage, such as cellular FLICE-like inhibitory proteins (cFLIPs), leads to the recruitment of receptor-interacting protein kinase 3 (RIPK3) and the subsequent formation of complex IIb (41). Alternatively, the formation of the complex IIb can also be initiated by interferon receptors (IFNRs), toll-like receptors (TLRs), and intracellular RNA- or DNA-sensing molecules (42). RIPK3 recruits the mixed lineage kinase domain-like pseudokinase (MLKL), the executioner kinase of necroptosis, to form the necrosome complex (complex IIc) (42). We previously reported that the inhibition of procaspase 8 cleavage during HCMV infection of monocytes is necessary for RIPK3 phosphorylation (30), although the co-stimulatory signal required to activate RIPK3 remains unknown. Regardless, MLKL is not activated despite the rapid activation of RIPK3, indicating that HCMV prevents the execution of necroptosis within HCMV-infected monocytes (30).

HCMV and murine cytomegalovirus (MCMV) encode for several inhibitors of the necroptotic pathway (31, 35). MCMV viral inhibitor of RIP activation (vIRA) is encoded by the viral *M45* gene and contains a RHIM domain preventing activation of RIPK1 and RIPK3 (43, 44). Although HCMV encodes a M45 homolog, UL45, it lacks the RHIM domain required for the interaction with RIPK1 or RIPK3 (45). Instead, the HCMV *UL36-* encoded vICA promotes the degradation of MLKL to suppress necroptosis, which is in addition to its caspase 8 inhibitory activity (46, 47). However, vICA is not expressed during quiescent infection of monocytes, indicating a distinct mechanism of necroptosis suppression (18, 19). Our previous study demonstrated that the inhibition of HCMV-induced autophagy within infected monocytes allows for the phosphorylation of MLKL and subsequently necroptosis (30). To date, the mechanism through which HCMV-induced autophagy blocks necroptosis within quiescently infected monocytes is unclear.

Here, we report that HCMV infection rapidly upregulates cFLIP long (cFLIP_L_) abundance to inhibit the cleavage of procaspase 8, as depletion of cFLIP_L_ within infected monocytes initiates the extrinsic apoptotic cascade while directing cell death away from the necroptotic pathway. cFLIP_L_ -mediated inhibition of caspase 8 and the simultaneous activation of TLR3, but not TNFR1 or TLR4, is required for RIPK3 activation following HCMV infection of monocytes. However, consistent with our previous studies, HCMV stimulation of RIPK3 activity does not phosphorylate and activate MLKL, which is dependent on the induction of autophagy following infection (30). We found HCMV-induced autophagy mediates the sequestration of MLKL within the nucleus of infected monocytes, preventing MLKL shuttling between the cytoplasm and nucleus. Importantly, the presence of a nuclear export inhibitor prevents necroptosis in HCMV-infected monocytes treated with an autophagy inhibitor, indicating sequestration of MLKL in the nucleus is critical to suppressing necroptosis. Overall, our study identifies key factors responsible for triggering of the necroptotic pathway within quiescently infected monocytes and provides insight into a novel mechanism through which HCMV-induced autophagy prevents to execution of necroptosis.

## Results

### HCMV infection increases FLIP_L_ abundance to block cleavage of procaspase 8

Circulating monocytes have a short lifespan of 48 hours (h) that can be accelerated in response to a viral infection (17–19). We reported that HCMV infection stimulates monocyte survival through the 48-h viability checkpoint by increasing the abundance of a select subset of antiapoptotic proteins capable of blocking the intrinsic apoptosis pathway (16, 22, 27, 48). Concurrently, HCMV impedes death receptor-mediated extrinsic apoptosis by preventing procaspase 8 cleavage (22, 30). However, the molecular mechanisms through which HCMV attenuates caspase 8 activation remain unclear. To elucidate how procaspase 8 cleavage is blocked during HCMV infection of monocytes, we first examined the protein levels of cellular FLICE-like inhibitory protein (cFLIP), a known cellular repressor of caspase 8. cFLIP primarily exists as 2 isoforms, cFLIP short (cFLIP_S_) and cFLIP long (cFLIP_L_), due to alternative splicing (41, 49). cFLIP_S_ is a potent inhibitor of caspase 8 by forming inactive heterodimers with procaspase 8 (41, 49). Surprisingly, we found little change in the abundance of cFLIP_S_ in monocytes following HCMV infection (Fig. 1A), suggesting a minimal role for cFLIP_S_ in preventing caspase 8 activation. cFLIP_L_ can act both in a pro- and an anti-apoptotic manner, where low levels promote cell death while high concentrations attenuate apoptosis (41, 49). We found HCMV infection induced a substantial increase in cFLIP_L_ abundance at 24 hours post-infection (hpi) that was sustained through 72 hpi (Fig. 1A, 1B). UV-inactivated HCMV particles (UV-HCMV) also increased cFLIP_L_ similar to “live” virus, suggesting viral entry is responsible for the elevated levels of cFLIP_L_ within infected monocytes (Fig. 1C, 1D). In support, treatment with soluble glycoprotein gB (sgB), but not soluble glycoprotein gH (sgH), was sufficient to increase cFLIP_L_ levels to those found in HCMV-infected monocytes (Fig. 1E, 1F). In line with our previously studies demonstrating the critical role of viral entry in stimulating a prosurvival state within infected monocytes (22, 27, 50–53), these data further show that gB-initiated signaling during viral entry increases cFLIP_L_ abundance.

**Figure 1.**
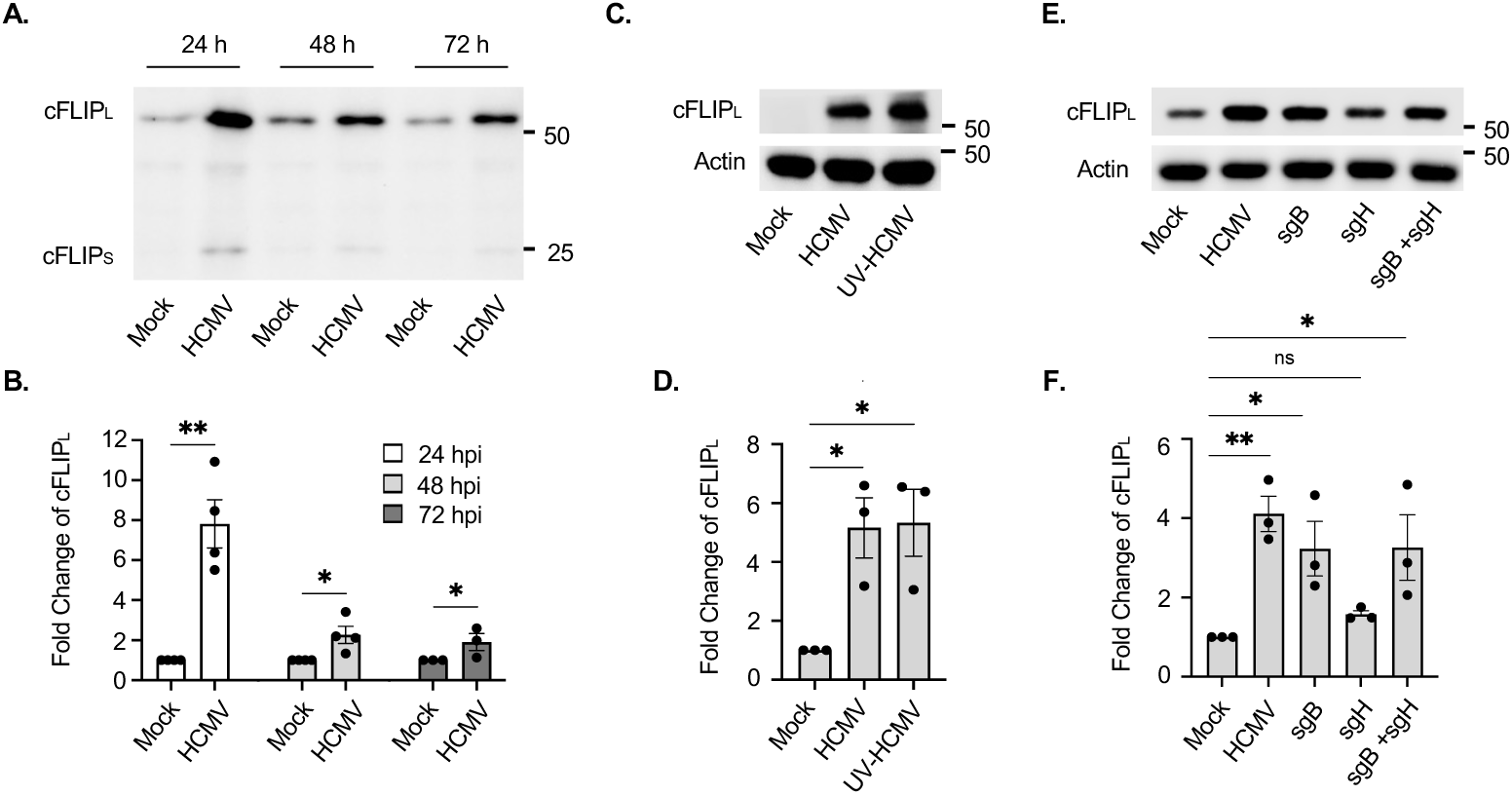
HCMV infection of monocytes increases cFLIP_L_ abundance. **(A to F)** Primary peripheral blood monocytes were mock infected or infected with HCMV (MOI 5) for 24 (A to F), 48 (A), or 72 h (A). **(E, F)** Cells were treated with 0.5 µg/mL of sgB, sgH, or both for 24 h. cFLIP_L_ was detected by Western blot (A, C, E) and fold change quantified (B, D, F). β-actin was used as a loading control. Western blots and densitometry are representative of at least 3 biological replicates per group. ns = not significant, *P<0.05, **P < 0.005, by one-way ANOVA with Tukey’s HSD post hoc test or Student’s t-test.

Next, we sought to determine if HCMV-induced cFLIP_L_ prevents the cleavage of procaspase 8 using a siRNA that targets all cFLIP isoforms (cFLIP siRNA #1) and a siRNA specific for cFLIP_L_ (cFLIP siRNA #2). Both siRNAs depleted cFLIP_L_ by ∼90% (Fig. 2). Consistent with our previous studies demonstrating HCMV prevents the intrinsic biological programming of monocytes to activate extrinsic apoptosis (22, 30), infection increased the levels of procaspase 8 with a corresponding decrease in the formation of the cleaved forms of caspase 8 (14 kDa and 18 kDa). The loss of cFLIP_L_ within infected monocytes reduced procaspase 8 while increasing caspase 8, suggesting the induction of cFLIP_L_ following HCMV infection prevents the initiation of the extrinsic apoptotic caspase cascade. Indeed, the activation of caspase 8 within cFLIP_L_-depleted, HCMV-infected monocytes led to the downstream cleavage of procaspase 3 into fully active caspase 3 (17 kDa). Thus, HCMV increases cFLIP_L_ levels to prevent cleavage of procaspase 8 and the subsequent initiation of the extrinsic apoptotic pathway.

**Figure 2.**
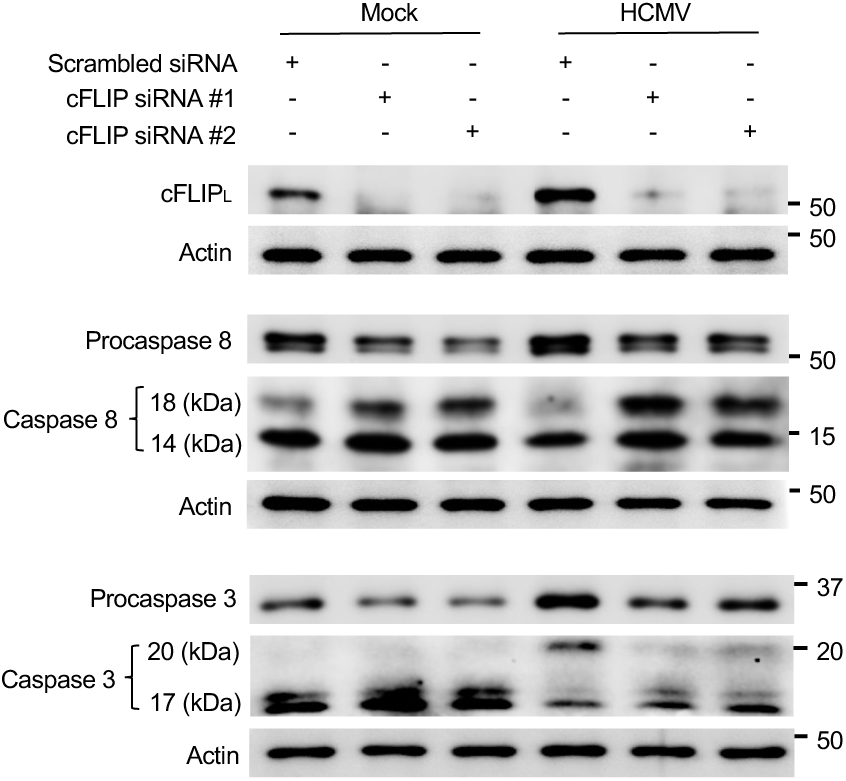
HCMV-induced cFLIP_L_ blocks cleavage of procaspase 8. Monocytes were mock infected or infected with HCMV (MOI 5) for 4 h. Following infection, cells were transfected with 500 nM of scrambled siRNA, cFLIP siRNA #1, or cFLIP siRNA #2 for 48 h. Total cFLIP, procaspase 8, caspase 8, procaspase 3, and caspase 3 were detected by Western blot. β-actin was used as a loading control. Western blots are representative of at least 3 biological replicates per group.

### TLR3 is required for HCMV-induced phosphorylation of RIPK3

Once cleavage of procaspase 8 is blocked, necroptosis is activated as a secondary cellular antiviral failsafe mechanism to promote the death of infected cells (54). As we have previously reported (30), HCMV infection increased both protein abundance and phosphorylation of RIPK3 (Fig. 3A, 3B), an essential component of the necrosome (41). Importantly, the release of procaspase 8 cleavage by siRNA-mediated depletion of cFLIP_L_ attenuated HCMV-induced RIPK3 abundance and phosphorylation, which is in line with other studies demonstrating cFLIP-mediated inhibition of procaspase 8 cleavage is required for opening the necroptotic “trapdoor” (34–37). Additionally, UV-HCMV infection alone increased RIPK3 levels and stimulated phosphorylation, indicating viral entry is sufficient to activate RIPK3 (Fig. 3C, 3D). However, treatment with sgB, sgH, or both was unable to stimulate protein levels or phosphorylation of RIPK3 (Fig. 3E, 3F) despite sgB increasing cFLIP_L_ abundance (Fig. 1E, 1F). These data suggest additional virally induced cellular signals independent of gB and gH are required to trigger necroptosis.

**Figure 3.**
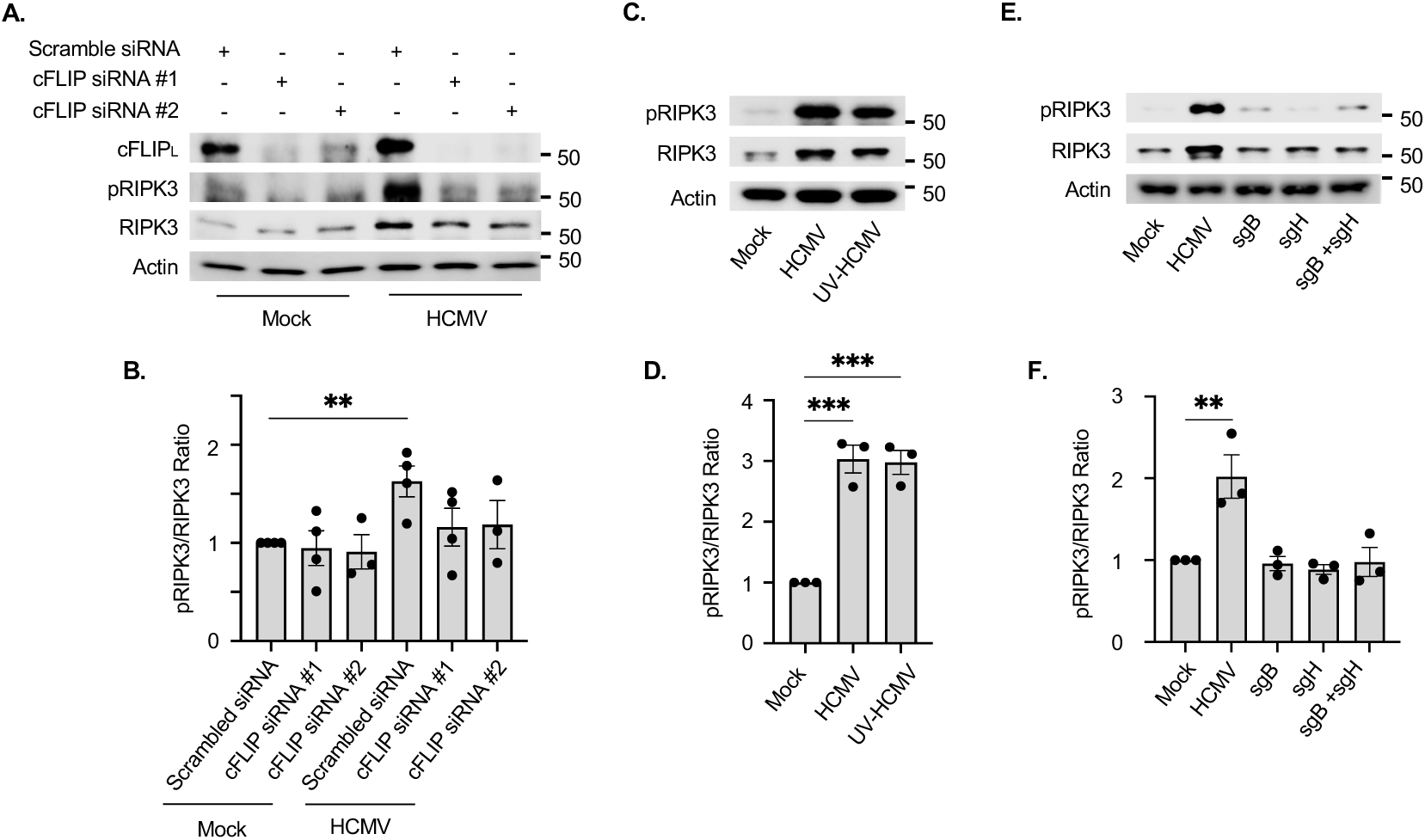
HCMV-induced cFLIP_L_ is required for RIPK3 phosphorylation. **(A, B)** Monocytes were mock infected or infected with HCMV (MOI 5) for 4 h. Following infection, cells were transfected with 500 nM of scrambled siRNA, cFLIP siRNA #1, or cFLIP siRNA #2 for 48 h. **(C to F)** Monocytes were mock infected or infected (MOI 5) with HCMV (C to F) or UV-HCMV (C, D) for 24 h. **(E, F)** Cells were treated with 0.5 µg/mL of sgB, sgH, or both for 24 h. cFLIP_L_, pRIPK3, and total RIPK3 were detected by Western blot (A, C, E). β-actin was used as a loading control. Western blots were quantified and the phosphorylation ratio of pRIPK3 to total RIPK3 was determined with scrambled siRNA or mock infected treatment groups set to 1 (B, D, F). Western blots and densitometry are representative of at least 3 biological replicates per group. **P < 0.005, *** P<0.005, by one-way ANOVA with Tukey’s HSD post hoc test or Student’s t-test.

Signaling from several receptors, including TNFR, TLR4, and TLR3, are known to promote RIPK3 phosphorylation in a cell-type-dependent manner (42). To identify if these receptors are involved in initiating the necroptotic pathway during HCMV infection of monocytes, we used a neutralizing antibody against TNFR (NAb TNFR1), the TLR4-selective inhibitor TAK-242 (iTLR4), or the TLR3 selective inhibitor (R)-2-(3-Chloro-6-fluorobenzo[b]thiophene-2-carboxamido)-3-phenylpropanoic acid (iTLR3). Activation of TNFR stimulates NF-κB signaling and necroptosis (55, 56). Accordingly, TNFα treatment induced phosphorylation of the inhibitor of nuclear factor kappa-B kinase subunit β (IKKβ) and RIPK3, which was abrogated by the presence of NAb TNFR1 (Fig. 4A). However, HCMV-induced RIPK3 phosphorylation was unaffected by the loss of TNFR1 signaling. LPS stimulates the phosphorylation of RIPK3, IRF3 (interferon signaling), and IKKβ in a TLR4-dependent manner. We found inhibition of TLR4 blocked RIPK3, IRF3, and IKKβ phosphorylation, but had little effect on HCMV-induced RIPK3 phosphorylation (Fig. 4B). In contrast, inhibition of TLR3 signal attenuated the phosphorylation of RIPK3 induced by both the cognate TLR3 ligand poly I:C and HCMV via a reduction of total protein levels (Fig. 4C), suggesting TLR3 signaling stimulates RIP3K abundance to increase the levels of phosphorylated RIPK3 (pRIPK3). In agreement, depletion of TLR3 by siRNA reduced the amount of pRIPK3 within HCMV-infected monocytes (Fig. 4D, 4E). To address if TLR3 also directly facilitates a rapid RIPK3 phosphorylation following HCMV infection, monocytes were pretreated with iTLR3 for 1 h and infected for 30 minutes (min) prior to any change in RIPK3 abundance (Fig. 4F, 4G). We found HCMV infection significantly increased in the ratio of pRIP3K to RIP3K, which was reduced by the presence of iTLR3 to levels comparable of control uninfected monocytes. Thus, TLR3 appears to respond to HCMV infection by increasing both RIPK3 protein abundance and phosphorylation levels in order to initiate the necroptotic pathway within infected monocytes.

**Figure 4.**
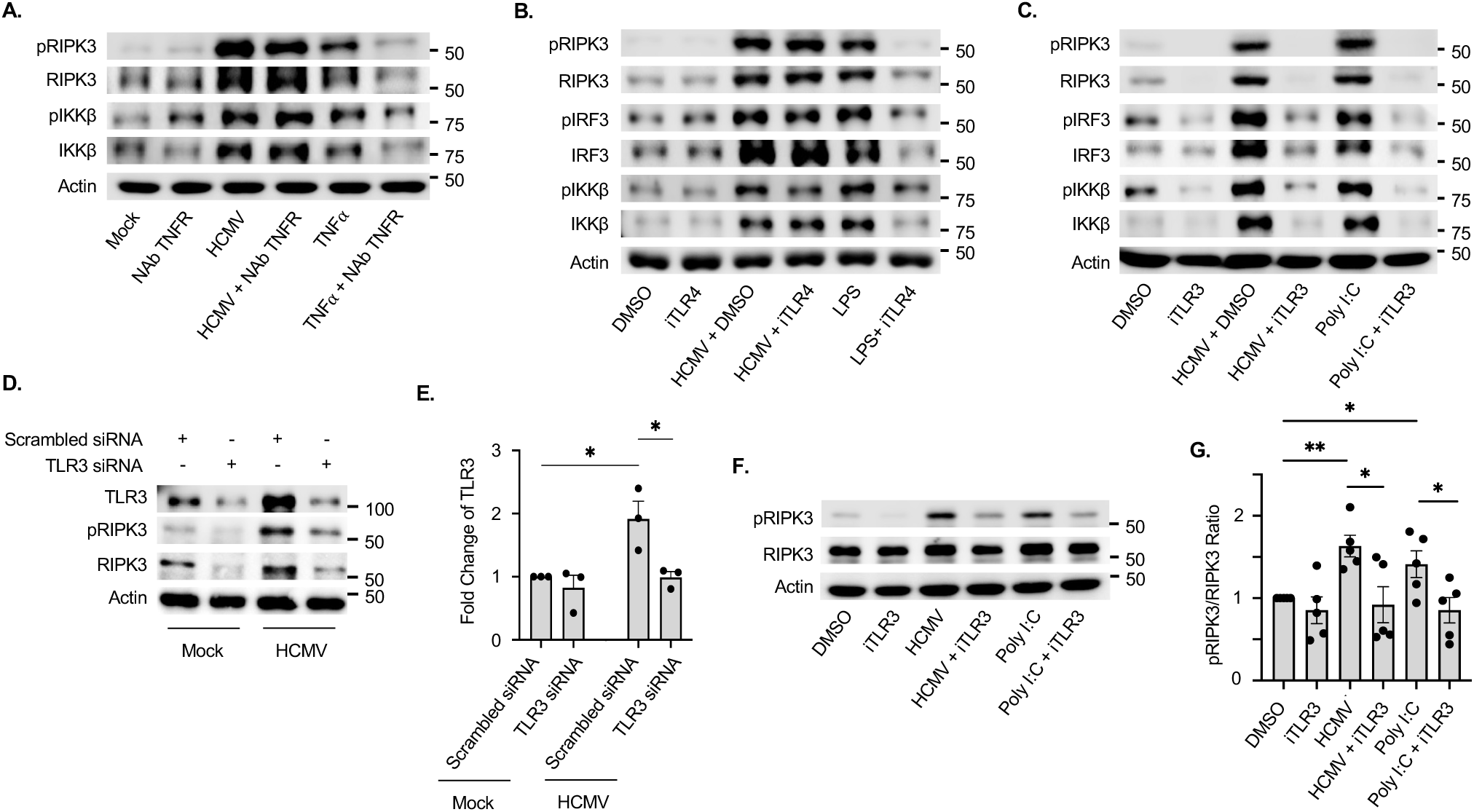
TLR3 is necessary the phosphorylation of RIPK3 following HCMV infection. **(A to C, F, G)** Monocytes were pretreated for 1 h with 1 µg/mL of TNFR neutralizing antibody (NAb TNFR), 1µM of TAK-242 (iTLR4; a TLR4 antagonist), or 100 nM of (R)-2-(3-Chloro-6-fluorobenzo[b]thiophene-2-carboxamido)-3-phenylpropanoic acid (iTLR3; a TLR3 antagonist). **(D, E)** Monocytes were transfected with 500 nM of a scrambled siRNA or a TLR3 siRNA for 48 h. Following pretreatment with inhibitors or depletion with siRNA, monocytes were mock infected or infected with HCMV (MOI 5) or treated with 2 nM of TNFα (a TNFR ligand), 0.1 ng/mL of LPS (a TLR4 ligand), or 1 µg/mL poly I:C (a TLR3 ligand) for 30 min (F, G) or 24 h (A to E). pRIPK3, total RIPK3, pIRF3, total IRF3, pIKKβ, and total IKKβ were detected by Western blot (A to D, F). β-actin was used as a loading control. Western blots were quantified and the fold change in TLR3 abundance determined with scrambled siRNA set to 1 (E) or the phosphorylation ratio of pRIPK3 to total RIPK3 determined with DMSO control group set to 1 (G). Western blots and densitometry are representative of at least 3 biological replicates per group. *P<0.05, **P < 0.005, by one-way ANOVA with Tukey’s HSD post hoc test or Student’s t-test.

### HCMV-induced autophagy blocks MLKL nucleocytoplasmic shuttling

HCMV infection initiates necroptosis signaling through the combined effects of cFLIP_L_ (Fig. 2) and TLR3 (Fig. 4). However, we previously reported that the induction of autophagy blocks the phosphorylation of MLKL and the execution of necroptosis within infected monocytes (30). To date, the mechanism through which autophagy prevents MLKL activation is unknown. Trafficking of MLKL between the cytoplasm and nucleus has been shown to be a key regulatory mechanism controlling MLKL activation (57–60). Thus, we examined the localization of MLKL within HCMV-infected monocytes in the presence or absence of Spautin-1 (SP-1), an autophagy inhibitor which we have previously validated to attenuate HCMV-induced autophagy (30). We found HCMV infection significantly increased the percent of MLKL (pink) localized to the nuclei (blue) of infected monocytes when compared to uninfected cells (Fig. 5A, 5B). The presence of SP-1 decreased to levels of nuclear MLKL to mock levels. Because we previously showed SP-1 had little effect on the total protein levels of MLKL within HCMV infected monocytes (30), our new data here suggests MLKL is not being degraded in the cytoplasm by autophagy but rather trapped within the nucleus of infected cells. In support, cytoplasmic and nuclear fractionation demonstrated increased MLKL levels in the nucleus of HCMV-infected monocytes, which was reduced by SP-1 treatment (Fig. 5C). Interestingly, the total levels of cytoplasmic MLKL remained constant between infected and uninfected monocytes, suggesting that the basal levels of total MLKL found in the cytoplasm are relatively unaffected by any changes in the rate of nucleocytoplasmic shuttling induced by HCMV. Examination of cytoplasmic phosphorylated MLKL (pMLKL) revealed that HCMV-infected monocytes express similar levels to mock-infected cells and that inhibition of autophagy by SP-1 increases cytoplasmic levels of pMLKL. The increase in cytoplasmic pMLKL corresponded to a decrease of total MLKL found in the nucleus of SP-1-treated, HCMV-infected monocytes. pMLKL was undetectable in the nucleus of infected cells, suggesting that exiting MLKL from the nucleus is being rapidly phosphorylated in the cytoplasm. To test if MLKL trafficking from the nucleus to the cytoplasm represents the pool of activated MLKL in the cytoplasm, we utilized a potent nuclear export inhibitor leptomycin B (LMB). LMB treatment alone had little effect on the nuclear abundance of total MLKL or the cytoplasmic levels of pMLKL in infected cells. However, LMB treatment of SP1-treated, HCMV-infected monocytes increased nuclear levels of total MLKL while reducing the cytoplasmic levels of pMLKL. Together, these data demonstrate that HCMV-induced autophagy suppresses MLKL from exiting the nucleus, thus preventing its cytoplasmic phosphorylation within infected monocytes.

**Figure 5.**
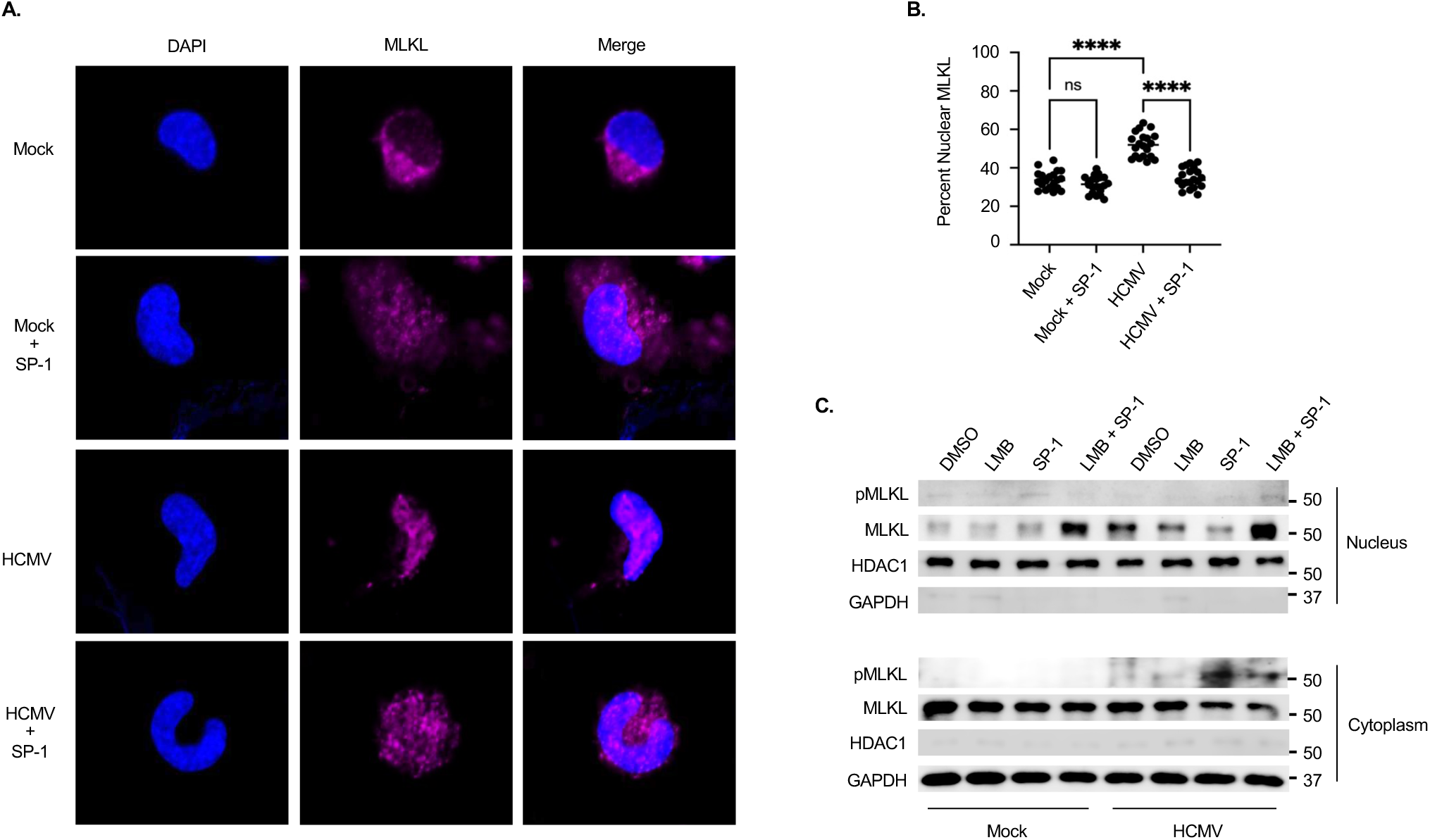
HCMV-induced autophagy sequesters MLKL in the nucleus of HCMV-infected monocytes. **(A to C)** Monocytes were pretreated with DMSO, 50 µM of spautin-1 (SP-1; an autophagy inhibitory), 1 nM leptomycin B (LMB; a nuclear export inhibitor), or both SP1 and LMB for 1-3 h. Cells were then mock infected or infected with HCMV (MOI 5) for 24 h. **(A)** Cells were stained for nuclei (DAPI; blue) and MLKL (pink). **(B)** Fiji plugin ComDet v0.5.5 was used to quantify MLKL cytoplasmic and nuclear fluorescence. Quantification of subcellular localization of MLKL was from at least 30 cells per biological replicate per group. **(C)** Subcellular fractionation was performed to isolate cytoplasmic and nuclear extracts. pMLKL and total MLKL were detected by Western blot. GAPDH and HDAC1 were used as cytosolic and nuclear loading controls. Immunofluorescent images and Western blots are representative of at least 3 biological replicates per group. ns = not significant, ****P<0.0001, by one-way ANOVA with Tukey’s HSD post hoc test or Student’s t-test.

### HCMV prevents necroptosis by blocking the nuclear export of MLKL

Next, we investigated if interrupting MLKL trafficking out of the nucleus of HCMV-infected monocytes is responsible for inhibiting necroptosis. Monocytes were mock- or HCMV-infected in the presence of SP-1, LMB, or both followed by staining with propidium iodide (PI) and annexin V (Fig. 6A), which allows for the differentiation between apoptotic and necroptotic/late death cells (30, 61). Consistent with our previous studies (22, 30, 50–53), HCMV infection increased cell survival (live gate; PI- and annexin V-) (Fig. 6B), as well as significantly reduced the rate of cells dying by apoptosis (PI- and annexin V+) (Fig. 6C), relative to uninfected cells. The inhibition of autophagy by SP-1 in HCMV-infected monocytes decreased cell viability (Fig. 6B) but had little effect on the frequency of apoptotic cells (Fig. 6C). In contrast, the presence of SP-1 significantly increased the rate of infected cells undergoing necroptosis (PI+ and annexin V+) (Fig. 6D), which we have previously shown to be reversed by the presence of a MLKL inhibitor (30). Strikingly, LMB treatment also reversed SP-1-induced necroptosis of HCMV-infected monocytes, as the percent of live cells (Fig. 6B) and those undergoing necroptosis (Fig. 6D) returned to levels observed in cells infected with HCMV alone, suggesting MLKL export from the nucleus of autophagy-inhibited, HCMV-infected monocytes is required for necroptosis. Thus, these data support a mechanism whereby HCMV-induced autophagy opposes the execution of necroptosis in infected monocytes occurs via retention of MLKL in the nucleus, thereby preventing its phosphorylation and subsequent pore forming activity.

**Figure 6.**
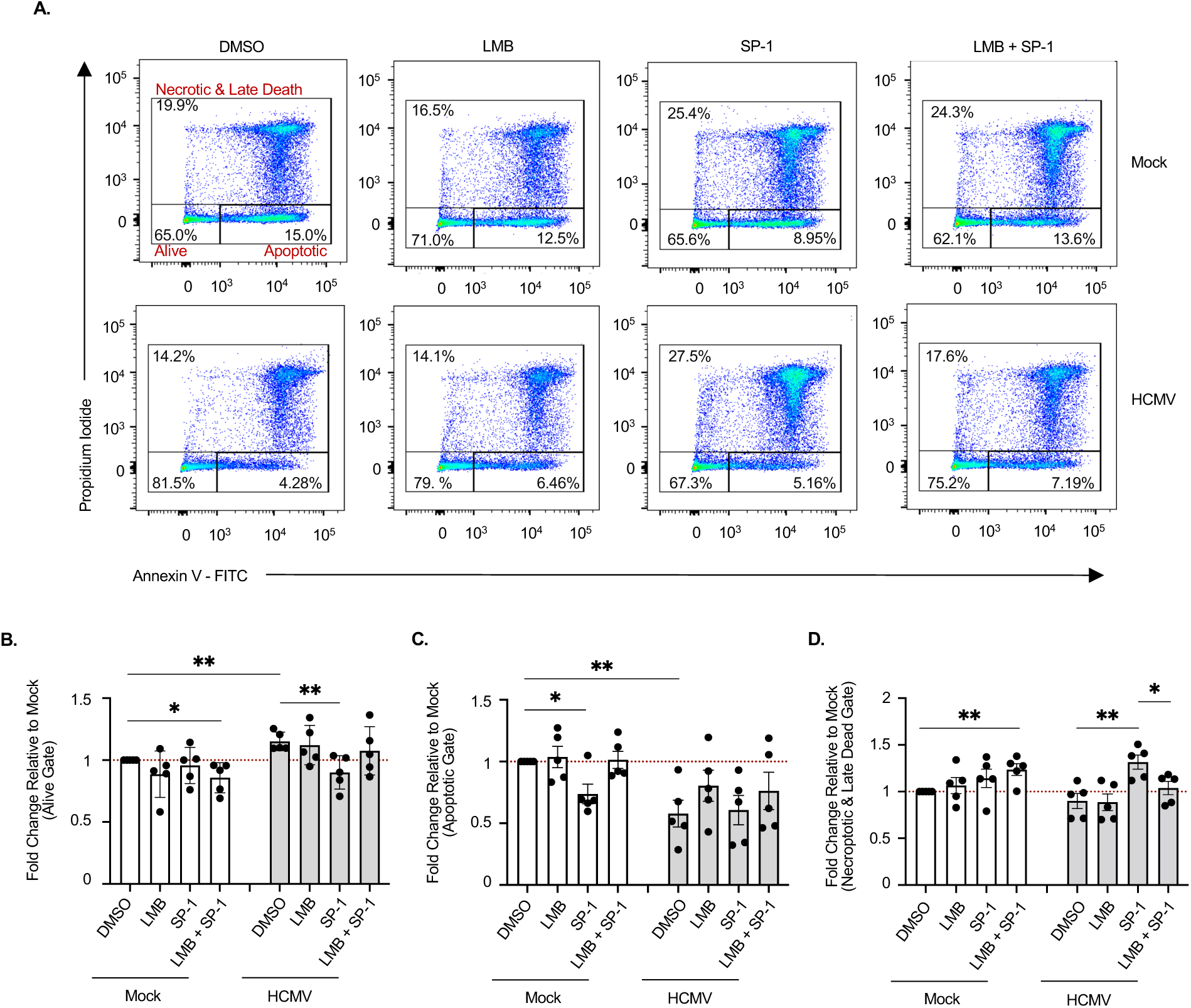
HCMV prevents export of nuclear MLKL to prevent necroptosis of infected monocytes. **(A to D)** Monocytes were pretreated with DMSO, 50 µM of SP-1, 1 nM of LMB, or both for 1-3 h. Cells were then mock infected or infected with HCMV (MOI 5) for 24 h. Cell viability was determined by annexin V and propidium iodide (PI) staining followed by flow cytometric analysis. All results are representative of at least 3 biological replicates per group. **P<0.005, *P<0.05, by one-way ANOVA with Tukey’s HSD post hoc test or Student’s t-test.

## Discussion

Peripheral blood monocytes are central players in mediating systemic dissemination of HCMV following a primary infection ultimately leading to lifelong persistence within the bone barrow of infected individuals (16, 62). However, monocytes are preprogrammed to undergo apoptosis ∼48 h after entry into the circulation from the bone marrow, which can be accelerated by cellular antiviral death responses (17, 63). Although HCMV has evolved a multitude of mechanisms to counteract apoptosis, the suppression of apoptosis can shift cell death pathways towards necroptosis as a secondary antiviral failsafe (30, 54, 64). We previously reported that the blockade of apoptosis induced by HCMV initiates necroptosis that was rapidly impeded by the viral induction of autophagy ensuring the survival of infected monocytes (30). To date, the mechanistic underpinnings responsible for 1) triggering necroptosis within HCMV-infected monocytes and 2) suppressing necroptosis following induction remain unknown. In this study, we demonstrate that HCMV upregulates cFLIP_L_ to prevent initiation of extrinsic apoptosis via the inhibition of procaspase 8 cleavage into active caspase 8 (Figs. 1, 2), a requirement for opening of the necroptotic trapdoor (54). A second signal from either a death receptor or a pathogen recognition receptor is then needed to initiate the necroptotic signaling cascade (39, 65). Although several receptors are known to trigger necroptosis, TLR3 is specifically required for the activation of RIPK3 within HCMV-infected monocytes (Fig. 4). Previous work for our lab determined HCMV concurrently induces autophagy to prevent activation of MLKL and stall the progression of necroptosis (30). We further show here that HCMV-induced autophagy disrupts cycling of MLKL between the cytoplasm and nucleus leading to the sequestration of MLKL within the nucleus of infected monocytes (Fig. 5). Overall, our study identifies novel viral countermeasure mechanisms designed to simultaneously oppose extrinsic apoptosis and necroptosis safeguarding the survival of infected monocytes.

Apoptosis can be categorized as intrinsic or extrinsic depending on the initiating signal (28, 66). We previously reported that HCMV upregulates a select subset of anti-apoptotic proteins, such as Mcl-1, HSP27, and XIAP, to block the intrinsic apoptotic pathway (22, 27, 53). Although, we also demonstrated that HCMV blocks the extrinsic apoptotic pathway by preventing procaspase 8 cleavage, the underlying mechanism remained undefined (22, 30). During lytic HCMV infection, HCMV *UL36*-encoded vICA directly binds and blocks procaspase 8 cleavage (67). However, the lack of vICA expression in quiescently infected monocytes suggests HCMV modulates the activity of cellular factors to suppress caspase 8 activation (18, 19). cFLIP is a critical regulator of caspase 8 activation in death receptor pathways and exists as two major isoforms, cFLIP_L_ and cFLIP_S_. cFLIP_S_ is a potent anti-apoptotic regulator that can directly bind and inhibit cleavage of procaspase 8 (41, 54, 68). Yet, HCMV has little effect on cFLIP_S_ abundance within infected monocytes (Fig. 1A), suggesting a minimal role in preventing the execution of extrinsic apoptosis. In contrast to cFLIP_S_, cFLIP_L_ contains catalytically inactive caspase-like domains in its C-terminal region and can have both pro- and anti-apoptotic activities, which are highly dependent on protein abundance (41, 54, 68). At low amounts, cFLIP_L_ forms a catalytically active heterodimer with procaspase 8 at the death-inducing signaling complex (DISC) of activated death receptors. Activation of procaspase 8 is achieved through stabilization of the active center of procaspase 8 by cFLIP_L_ without the requirement of proteolytic cleavage (29, 54). At high concentrations, cFLIP_L_ inhibits apoptosis due to the replacement of procaspase 8 at the DISC (41, 49). Here, we demonstrate HCMV profoundly affects the ratio of cFLIP_L_ to procaspase 8 by stimulating a robust increase of cFLIP_L_ within infected monocytes (Fig. 1). This stochiometric shift towards cFLIP_L_ likely impedes the recruitment of procaspase 8 to the DISC of death receptors, which is consistent with our new data showing HCMV-induced cFLIP_L_ is necessary for preventing caspase 8 activation and the subsequent initiation of the extrinsic apoptosis pathway following infection.

Inhibition of caspase 8 activation during viral infections triggers necroptosis as a secondary antiviral cell death response pathway. Given the importance of cFLIP in the regulation of caspase 8 activity, high levels of cFLIP_L_ promote the assembly of the necrosome (RIPK1, RIPK3, MLKL, FADD, and procaspase 8) ultimately leading to RIPK3 auto-phosphorylation and MLKL activation (41, 49). In agreement, depletion of cFLIP_L_ in HCMV-infected monocytes prevents the activation of RIPK3 (Fig. 3) shifting cell death back towards the apoptotic pathway (Fig. 2). In addition to the inhibition of caspase 8 activity, necroptosis requires an initiation signal typically originating from either a death receptor or a pathogen recognition receptor (34–37). Although our previous work demonstrated HCMV activates RIPK3, the specific trigger responsible for initiating the necroptotic pathway has remained elusive. Our new data here shows activation of TLR3 is specifically responsible for increasing the levels of phosphorylated RIPK3 within HCMV-infected monocytes via directly stimulating phosphorylation and increasing total protein abundance (Fig. 5). How TLR3 signaling is initiated during HCMV infection is unclear. HCMV has been shown to activate TLR2 signaling through direct binding with gB (69). To date, a TLR3-binding HCMV glycoprotein has yet to be identified, which is consistent with neither sgB nor sgH being able to increase the levels of phosphorylated RIPK3 (Fig. 3E, 3F). Alternatively, TLR3 canonically recognizes extracellular dsRNA delivered by incoming viruses during entry or produced during viral gene expression (70). Herpesviruses, including HSV-1, EBV, KSHV, and HCMV, contain viral RNAs within the tegument that could activate TLR3 signaling (71–74). Consistent with this possibility, UV-HCMV stimulates RIPK3 phosphorylation and increases protein abundance similar to replication competent virus (Fig. 3C, 3D). Regardless of the mechanism of TLR3 activation, our study demonstrates that the concomitant increase in cFLIP_L_ levels and signaling from TLR3 are necessary to trigger the early steps of the necroptotic cascade following HCMV infection of monocytes.

HCMV infection rapidly stimulates autophagy to block the execution of necroptosis by preventing activated RIPK3 from phosphorylating MLKL (30). The mechanism by which HCMV-induced autophagy blocks this critical step of the necroptotic pathway within infected monocytes was unclear, although we showed autophagy to attentuate the interaction between RIPK3 and MLKL (30). Recent studies demonstrated that the nucleocytoplasmic shuttling of MLKL can be a critical regulatory step in controlling MLKL-mediated necroptosis (59, 60). Here, we found HCMV infection traps unphosphorylated MLKL within the nucleus of infected monocytes to prevent necroptosis (Fig. 5), which is consistent with a previous report demonstrating the pharmacological inhibition of the nuclear export machinery leads to the accumulation of MLKL in the nucleus and a reduction in cell death (59). Since infected monocytes appear to contain high basal levels of unphosphorylated cytoplasmic MLKL (Fig. 5C), our data further suggest that only MLKL cycling out of the nucleus of infected monocytes is licensed to be phosphorylated when autophagy is inhibited. Nuclear autophagy is an emerging field that has been implicated in playing a key role in nuclear export and autophagic substrate encapsulation (75, 76). LC3B, a central protein in autophagy, associates with the nuclear membrane to form nuclear autophagosomes (75, 77), and SIRT1, a nuclear deacetylase, mediates LC3B activation (78). HCMV infection of monocytes increases the abundance and activity of both LC3B and SIRT1 (30, 79), perhaps hinting at the involvement of nuclear phagophore formation in sequestering MLKL within the nucleus. Regardless, our study identifies a novel nuclear retention mechanism of MLKL employed by HCMV to suppress necroptosis in quiescently infected monocytes.

In summary, our study demonstrates HCMV inhibits extrinsic death receptor-mediated apoptosis by increasing the abundance of cFLIP_L_ following infection of monocytes. Blockade of procaspase 8 activation by cFLIP_L_ results in the opening of the cellular trapdoor antiviral death pathway necroptosis, which is then triggered through the recognition of HCMV infection by TLR3. To combat necroptosis, HCMV stimulates autophagy to sequester MLKL in the nucleus of infected monocytes thereby preventing the execution of necroptosis. Overall, our study highlights the complex “arms race” between host antiviral cellular death pathways and HCMV countermeasures designed to thwart these responses, thus allowing monocytes to act as vehicles for viral dissemination. Elucidating the multitude of mechanisms responsible for ensuring the survival of quiescently infected monocytes may shed light on novel host-directed antivirals.

## Materials and Methods

### Human peripheral blood monocyte isolation and culture

Isolation of human peripheral blood monocytes was performed as previously described (27, 80). Briefly, blood was drawn from random deidentified donors by venipuncture, diluted in Roswell Park Memorial Institute medium (RPMI) 1640 (ATCC, Product #30-2001), and centrifuged through Histopaque 1077 (MilliporeSigma) to remove red blood cells and neutrophils. Mononuclear cells were collected and washed with saline to remove the platelets and then separated by centrifugation through a Percoll (GE Healthcare) gradient (40.48% and 47.70%). More than 90% of isolated peripheral blood mononuclear cells were monocytes as determined by CD14- or CD16-postive staining. Cells were washed with saline, resuspended in RPMI 1640 and counted. All experiments were performed in the absence of human serum (unless mentioned otherwise) at 37°C in a 5% CO_2_ incubator. SUNY Upstate Medical University’s Institutional Review Board and Health Insurance Portability and Accountability Act guidelines for the use of human subjects were followed for all experimental protocols in our study (IRB#: 262458-19). For the inhibitor studies, the following reagents were used: Spautin-1 (SP-1; a USP10 and USP13 inhibitor), SBI-0206965 (SBI; a ULK1 inhibitor), TAK-242 (iTLR4; a TLR4 inhibitor) from Selleckchem. TNFR neutralizing antibody (NAb TNFR) from R&D systems. (R)-2-(3-Chloro-6-fluorobenzo[b]thiophene-2-carboxamido)-3-phenylpropanoic acid (iTLR3; a TLR3 inhibitor) from MilliporeSigma. LPS and TNFα from Invitrogen. Poly I:C from TOCRIS (Minneapolis, MI).

### Virus preparation and infection

All virus stocks were propagated on human embryonic lung (HEL) 299 fibroblasts (CCL-137, ATCC) of low passage (P7-15) in Dulbecco’s Modified Eagle medium (DMEM) (Lonza) with 2.5 μg/ml plasmocin (Invivogen) and 10% fetal bovine serum (FBS) (MilliporeSigma). When a 100% cytopathic effect was observed, virus was purified from the supernatant by ultracentrifugation (115000 x *g*, 65 minutes (min), 22°C) through a 20% sorbitol cushion to remove cellular contaminants and resuspended in RPMI 1640 medium (ATCC, Product # 30-2001). A multiplicity of infection (MOI) of 1 genome copy/cell was used for each experiment unless otherwise stated. UV-inactivated virus was prepared by incubating virus in a BioRad GS Linker UV Chamber (UV wavelength, 254 nm) for 360s on ice. All UV-inactivated virus preparations were confirmed to not produce any detectable levels of de novo synthesized viral gene products.

### Western blot analysis

Monocytes were harvested in a modified radioimmunoprecipitation assay (RIPA) buffer (50 mM Tris-HCl [pH 7.5], 5 mM EDTA, 100 mM NaCl, 1% Triton X-100, 0.1% SDS, 10% glycerol) supplemented with protease inhibitor cocktail (Sigma-Aldrich St. Louis, MO) and phosphatase inhibitor cocktails 2 and 3 (Sigma-Aldrich St. Louis, MO) for 30 minutes (min) on ice. The lysates were cleared from the cell debris by centrifugation at 4°C (5 min, 21000 × g) and stored at - 20°C until further analysis. Protein samples were solubilized in Laemmli SDS-sample nonreducing (6×) buffer (Boston Bioproducts) supplemented with β-mercaptoethanol (Amresco) by incubating at 95°C for 10 min. Equal amounts of total protein from each sample were loaded in each well, separated by SDS-polyacrylamide gel electrophoresis, and transferred to polyvinylidene difluoride membranes (Bio-Rad). Blots were blocked in 5% bovine serum albumin (BSA; Fisher Scientific) for 1 h at room temperature and then incubated with primary antibodies overnight at 4°C. The following antibodies were used: anti-FLIP, anti-procaspase 8, anti-p-RIPK3 (S227), anti-pIRF3 (Ser396), anti-IRF3, anti-pIKKα/β (Ser176/180), anti-IKKβ, anti-TLR3, anti-MLKL, anti-p-MLKL (S358), anti-pAKT (S473), anti-AKT, anti-HDAC1 and anti-GAPDH were from Cell Signaling & Technology; anti-caspase 3 and anti-RIPK3 antibodies were from Santa Cruz; anti-IE1 antibody was a generous gift from Tom Shenk (81); rhodamine anti-actin antibody was from Bio-Rad. Blots were then incubated with: 1) horseradish peroxidase (HRP)-conjugated secondary antibodies (Cell Signaling) for 30 min at room temperature and chemiluminescence was detected using the Clarity Western ECL substrate (Bio-Rad) or 2) alkaline phosphatase (AP) conjugated secondary antibodies (Promega) for 1 h at RT and colorimetric detection performed using AP Conjugate Substrate Kit (Bio-Rad). Densitometry was performed using Image Lab software (Bio-Rad).

### Cellular Fractionation Lysis Gradient

Subcellular fractionation was performed with an iso-osmotic discontinuous iodixanol-based density gradient as previously described with minor modifications (82, 83). Briefly, live monocytes (4×10^6^ monocytes) were loaded on top of an iso-osmolar discontinuous iodixanol (MilliporeSigma) based gradient. Cells were centrifuged at 1000 x *g* for 10 min (minutes) in a swinging bucket rotor. During centrifugation, monocytes travel through a preliminary cell wash layer prior to encountering a mild cell lysis layer (0.5% IPEGAL CA-630) (MilliporeSigma), which disrupts the plasma membrane while leaving nuclei intact. Undamaged nuclei then pass through a subsequent wash layer prior to encountering a hyper-dense float layer. Soluble cytoplasmic fractions were isolated from the cell lysis layer and crude nuclei were harvested from the interface between the second wash and float layer. Both cytoplasmic and nuclear fractions were prepared for Western blot analysis.

### siRNA Silencing

Primary monocytes (3 × 10^6^ cells/transfection) were washed with phosphate-buffered saline (PBS) and resuspended in 100 μl of P3 Primary Cell Nucleofector solution (Lonza) containing either a TLR3-specific Silencer Select siRNA (1μM) (Ambion-Thermo Fisher Scientific), a cFLIP-specific Silencer Select siRNA (500 nM) (Ambion-Thermo Fisher Scientific), a cFLIP_L_-specific siRNA (500 nM) (Dharmacon, 5’ AAGGAACAGCUUGGCGUUCAAUU 3’), or a Silencer negative-control siRNA (Ambion-Thermo Fisher Scientific). Transfection was performed with a 4D-Nucleofector system (Lonza) using program EI-100. Following transfection, monocytes were incubated in RPMI 1640 supplemented with 2% human AB serum at 37°C and allowed to recover for 24 h. Monocytes were then mock infected or infected with HCMV for 24 h and subjected to Western blot analysis.

### Purification of soluble sgB and sgH from stably expressing Expi293F cells

Soluble gB (sgB) and gH (sgH) glycoproteins were purified from stable sgB or sgH Expi293F cell lines, which we have previously established and described in (51). For isolation of soluble glycoproteins, stable expression cell lines were grown in Expi293 expression media (ThermoFisher Scientific) with 200 µg/ml geneticin at 8% CO_2_ on an orbital shaker (125 rpm). Following cell lysis, recombinant sgB or sgH were purified using Ni-charged resin (Bio-Rad) and dialyzed with PBS at the final stage of purification. Monocytes were treated with the soluble glycoproteins at 1 µg/ ml for each experiment, unless otherwise stated. Ni-charged resin purified lysate of un-transfected Expi293 cells was dialyzed with PBS and same amount of total volume of soluble glycoproteins was used as negative control.

### Immunofluorescence

Monocytes were fixed for 15 min in 4% paraformaldehyde (PFA) followed by washing with PBS twice. Cell permeabilization and blocking of nonspecific binding were performed by incubating the cells with 0.1% Triton X-100, 5% BSA, and human FcR blocking reagent (Miltenyi) in PBS for 30 min at room temperature. Cells were then incubated overnight at 4°C in a humidified chamber with an anti-MLKL antibody (Cell Signaling). Monocytes were then washed in PBS and incubated with an anti-mouse antibody conjugated to Alexa Fluor 647 and further incubated overnight at 4°C. Coverslips were mounted with ProLong Gold Antifade with DAPI (4′,6-diamidino-2-phenylindole) (Thermo Fisher). Stained cells were visualized with a 63x objective using a Marianas system (3i) microscope (Okolab). The acquisition is done using SlideBook 6 (3i) and exported into Fiji/ImageJ software (an open Java-based image processing program developed at the National Institutes of Health for analysis and quantification.

### Statistical Analysis

All experiments were performed with a minimum of 3 biological replicates per group. Data were analyzed using one-way ANOVA with Tukey’s honest significant difference (HSD) post hoc test with GraphPad Prism software, and *p*-values less than 0.05 were considered statistically significant.

## Acknowledgements

We thank Chris Burrer in the Department of Microbiology and Immunology at SUNY Upstate Medical University for technical support, maintenance of lab operations, and assistance with virus growth and isolation. This work was supported by grants from the National Institute of Allergy and Infectious Disease (R01AI170834 and R01AI141460) to G.C. Chan, and National Heart, Lung, and Blood Institute (R01HL139824) to G.C. Chan.

